# Pangenomics reveal diversification of enzyme families and niche specialization in globally abundant SAR202 bacteria

**DOI:** 10.1101/692848

**Authors:** Jimmy H.W. Saw, Takuro Nunoura, Miho Hirai, Yoshihiro Takaki, Rachel Parsons, Michelle Michelsen, Krista Longnecker, Elizabeth B. Kujawinski, Ramunas Stepanauskas, Zachary Landry, Craig A. Carlson, Stephen J. Giovannoni

## Abstract

It has been hypothesized that abundant heterotrophic ocean bacterioplankton in the SAR202 clade of the phylum *Chloroflexi* evolved specialized metabolism for the oxidation of organic compounds that are resistant to microbial degradation via common metabolic pathways. Expansions of paralogous enzymes were reported and implicated in hypothetical metabolism involving monooxygenase and dioxygenase enzymes. In the metabolic schemes proposed, the paralogs serve the purpose of diversifying the range of organic molecules that cells can utilize. To further explore this question, we reconstructed SAR202 single amplified genomes and metagenome-assembled genomes from locations around the world, including the deepest ocean trenches. In analyses of 122 SAR202 genomes that included six subclades spanning SAR202 diversity, we observed additional evidence of paralog expansions that correlated with evolutionary history, and further evidence of metabolic specialization. Consistent with previous reports, families of flavin-dependent monooxygenases were observed mainly in the Group III SAR202, in the proposed class *Monstramaria* and expansions of dioxygenase enzymes were prevalent in Group IV. We found that Group I SAR202 encode expansions of racemases in the enolase superfamily, which we propose evolved for the degradation of compounds that resist biological oxidation because of chiral complexity. Supporting the conclusion that the paralog expansions indicate metabolic specialization, fragment recruitment and fluorescence *in situ* hybridization with phylogenetic probes showed that SAR202 subclades are indigenous to different ocean depths and geographical regions. Surprisingly, some of the subclades were abundant in surface waters and contained rhodopsin genes, altering our understanding of the ecological role of SAR202 in stratified water columns.

**Importance:** The oceans contain an estimated 662 Pg C of dissolved organic carbon (DOC). Information about microbial interactions with this vast resource is limited, despite broad recognition that DOM turnover has a major impact on the global carbon cycle. To explain patterns in the genomes of marine bacteria we propose hypothetical metabolic pathways for the oxidation of organic molecules that are resistant to oxidation via common pathways. The hypothetical schemes we propose suggest new metabolism and classes of compounds that could be important for understanding of the distribution of organic carbon throughout the biosphere. These genome-based schemes will remain hypothetical until evidence from experimental cell biology can be gathered to test them, but until then they provide a perspective that directs our attention to the biochemistry of resistant DOM metabolism. Our findings also fundamentally change our understanding of the ecology of SAR202, showing that metabolically diverse variants of these cells occupy niches spanning all depths, and are not relegated to the dark ocean.

## Introduction

Some dissolved organic matter (DOM) consists of labile molecules (LDOM) that are recycled quickly by microbes in the epipelagic (0-200 m) near the point of origin, while other DOM transits marine food webs and eventually accumulates in the deep ocean in the form of refractory dissolved organic matter (RDOM). RDOM has residence times of thousands of years (2) and is distributed throughout the water column, but is the main DOM type in the bathypelagic realm (>1000 m). Here we use the term *semi-labile DOM* (SLDOM) to encompass molecules that span a broad range of intermediate stabilities in the environment, including compounds that are often referred to as *recalcitrant* (3). Two general hypotheses put forward to explain SLDOM and RDOM are the *intrinsic stability hypothesis*, which postulates that DOM stability is due to molecular structures that are resistant to enzymatic cleavage (8), and the *molecular diversity hypothesis*, which predicts that extreme dilution of compounds can render them unusable by heterotrophs (4). Here, in genomes of the SAR202 clade of marine bacteria, we explore metabolic diversity related to both the *intrinsic stability hypothesis* and the *molecular diversity hypothesis*.

The first reports on SAR202 used molecular data to demonstrate their relative abundance increases dramatically at the transition between the euphotic and aphotic zones of the oceans (5). Microbes adapted to dark ocean regions (mesopelagic, 200-1000 m; bathypelagic, 1000-4000 m; abyssopelagic, 4000-6000 m; hadalpelagic, 6000-11,000 m) exploit environments where the most abundant energy resources are SLDOM. These compounds mainly are remnants from primary production in the epipelagic, which is attenuated in transit through food webs. In the dark oceans, low levels of primary production also occur locally, fueled by chemoautotrophy (6). The Microbial Carbon Pump (MCP) is a conceptual framework that captures these features of food webs, and recognizes that, in the process of transformation, a fraction of labile DOM is chemically altered to forms that resist or escape microbial degradation (7).

SAR202 are the most abundant lineage of bacteria in the deep oceans. This clade diversified approximately 2 billion years ago, forming six subclades, referred to as “Groups I-VI”) (9, 10). Early work showed that they constitute, on average, about 10% of total bacterioplankton throughout the mesopelagic of the Sargasso Sea, Central Pacific Ocean, and Eastern Pacific coastal waters (11). A subsequent study revealed that they constitute up to 5% of the total bacterioplankton community in the epipelagic and up to 30% in the meso- and bathypelagic zones in parts of the Atlantic Ocean (12).

SAR202 have escaped cultivation to date. Insight into their metabolism has come from field studies and comparative genomics (13). Recent studies, using both single-cell and metagenomic sequencing, have highlighted the differing roles for SAR202 groups at sites around the world. One study assembled three nearly complete SAR202 MAGs from metagenomes from oxygen minimum zones in the Gulf of Mexico and observed expression of nitrate reductase genes, suggesting these cells have the capacity for anaerobic respiration (14). Another study investigated vertical stratification and concluded that SAR202 might be sulfite oxidizers that utilize organosulfur compounds (15). An investigation of SAR202 from the Arctic Ocean described expanded families of dioxygenase enzymes that were proposed to function in aromatic compound degradation, potentially utilizing organic matter discharged from terrestrial sources (16). Freshwater relatives of SAR202 have also been discovered, shedding light on their diversity and ecology in aquatic habitats (17).

In a recent study of Group III SAR202, we identified expansions of paralogous protein families, including powerful oxidative enzymes that we hypothesized play a role in degrading SLDOM (10). SAR202 flavin-dependent monooxygenases (FMNOs) were hypothesized to oxidize a variety of chemically stable SLDOM molecules by introducing single oxygen atoms, for example by oxidizing sterols and hopanoids to carboxyl-rich alicyclic molecules (CRAM) (10). CRAM consists of fused aromatic and heterocyclic rings decorated with carboxyl groups (18–20).

In this study we investigated paralogous gene expansions and gene co-occurrence in a larger sample of SAR202 diversity. We reconstructed 10 new SAGs, isolated from mesopelagic and hadal waters from the Northwestern Pacific Ocean, and 73 new MAGs from the Bermuda Atlantic Time-series Study (BATS) site in the Sargasso Sea, and from TARA Oceans Expedition metagenomes, a total of 83 new SAR202 genomes. We also investigated the biogeography of these genomes, and their distribution as a function of depth in water columns. Interpreting this information, we hypothesize that SAR202 evolved and diversified into multiple niches where they play roles in the oxidation of resistant classes of DOM.

## Results

### Overview of genomic bins and SAGs

The total number of SAGs and MAGs in this study was 122, of which 83 are new, and the remainder from previous studies (10, 14, 21–23). Ten new SAR202 SAGs were obtained from three deep ocean trench stations: Mariana, Ogasawara, and Japan trenches. Sixty-two new SAR202 MAGs were reconstructed from TARA Oceans metagenome re-assemblies in this study. TARA metagenomic samples from different depths were assembled separately to help us preserve depth information for each MAG. Eleven new SAR202 MAGs came from metagenomic samples obtained at Bermuda Atlantic Time-series Study (BATS) site. A table summarizing the origin and depth of samples from which the SAGs and MAGs were obtained is provided as Supplemental Table 1.

### SAR202 diversity revealed through phylogenomic analyses

A phylogenomic tree was constructed from 36 concatenated single-copy genes that were selected based on their broad presence in genomes, suggesting core functions, and evidence of linear inheritance (Fig. 1). Using ChloNOG subset of gene clusters from the eggNOG database, we identified 639 orthologous gene clusters that are present as single copies in 141 genomes (122 SAR202, 17 other *Chloroflexi*, and 2 cyanobacteria outgroup).

**Figure 1:**
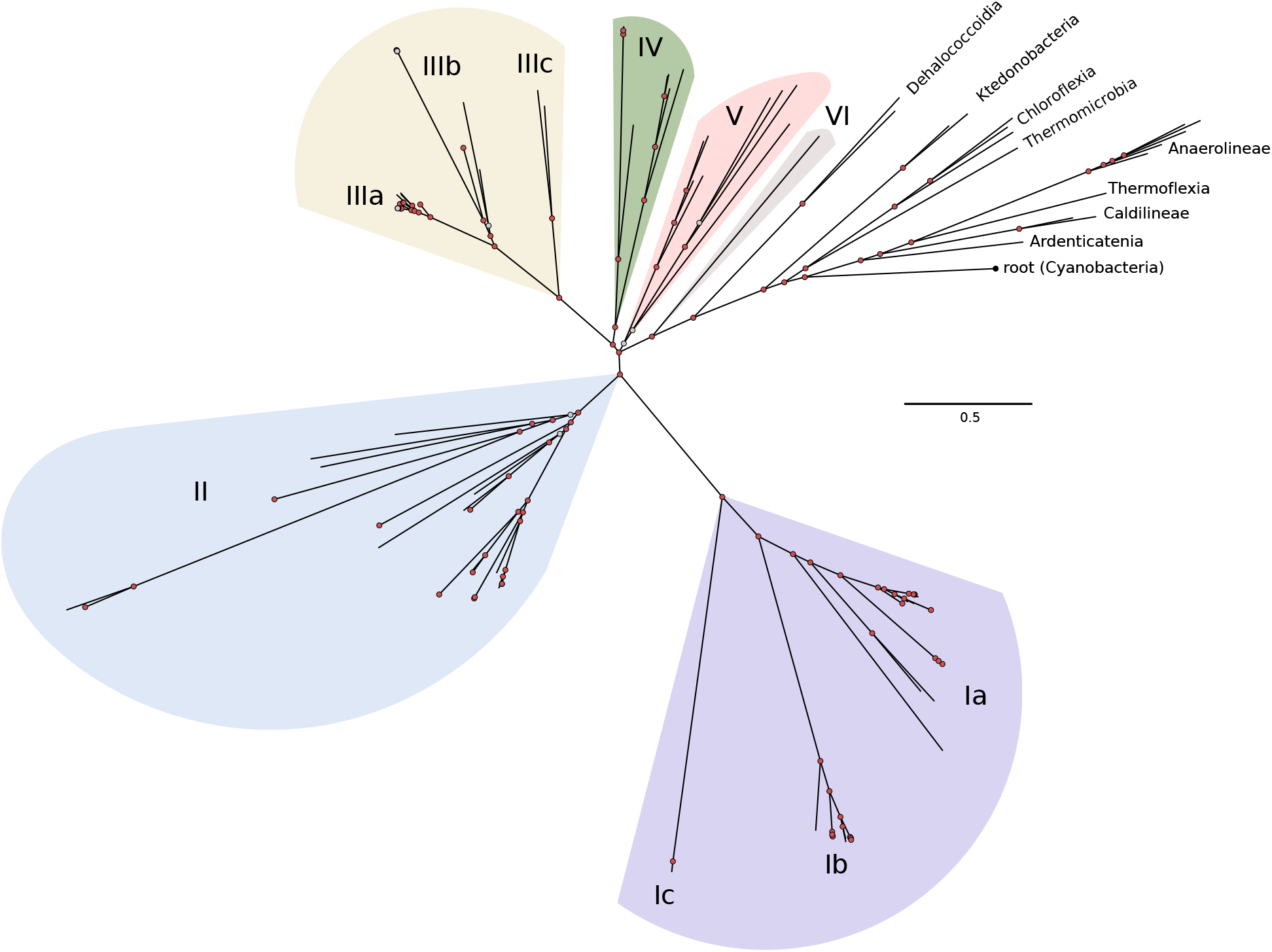
Phylogenomic tree of SAR202 genomes, built using 36 concatenated chloNOGs. Phylogenomic inference was done using Phylobayes MPI version 1.7. Cyanobacterial sequences were used for the outgroup. Color shading identifies SAR202 groups used in subsequent figures.

The phylogenomic tree supported earlier findings showing that SAR202 are a deeply-branching monophyletic group that radiates from within the *Chloroflexi*, possibly associated with *Dehalococcoides* (Fig. 1). Several deeply-branching subclades, Groups IV-VI, radiate near the base of the clade. Groups III, II and I appear in that order, ascending from the root. They are separated by large evolutionary distances and are the most abundantly represented SAR202 subgroups (Supplemental Table 1). Previously, we proposed that Group III be given the rank of class and assigned the name *Candidatus* Monstramaria (classis nov.). Given the separation of the subclades and the evolutionary distances between them in the phylogenomic tree, we propose the following names for the rest of SAR202 groups: Group I (*Candidatus* Umibozia, classis nov.), Group II (*Candidatus* Scyllia, classis nov.), Group IV (*Candidatus* Makaraia classis nov.), Group V (*Candidatus* Cetusia, classis nov.), and Group VI (*Candidatus* Tiamatia, classis nov.).

### Overview of paralogous enzyme superfamilies in SAR202

Paralog expansions, especially diverse, ancient ones, can indicate past evolutionary events in which new enzyme activities were vehicles for niche expansions. Investigating paralog expansions across SAR202 genomes, we constructed a heatmap showing relative abundances of the top 50 most abundant COG categories (Fig. 2A). The heatmap revealed five major expansions of paralogous gene families, and many other less prominent expansions. The distributions of these groups of paralogs across the major SAR202 subclades are shown in Fig. 2B. COG4948, the enolase superfamily, were mainly found in Group I and Group II (Fig. 2B); COG2141, the SAR202 FMNO paralogs were found mainly in Group II and III; and COG4638, ring-hydroxylating dioxygenase paralogs, were found in Group IV, as reported previously (16).

**Figure 2:**
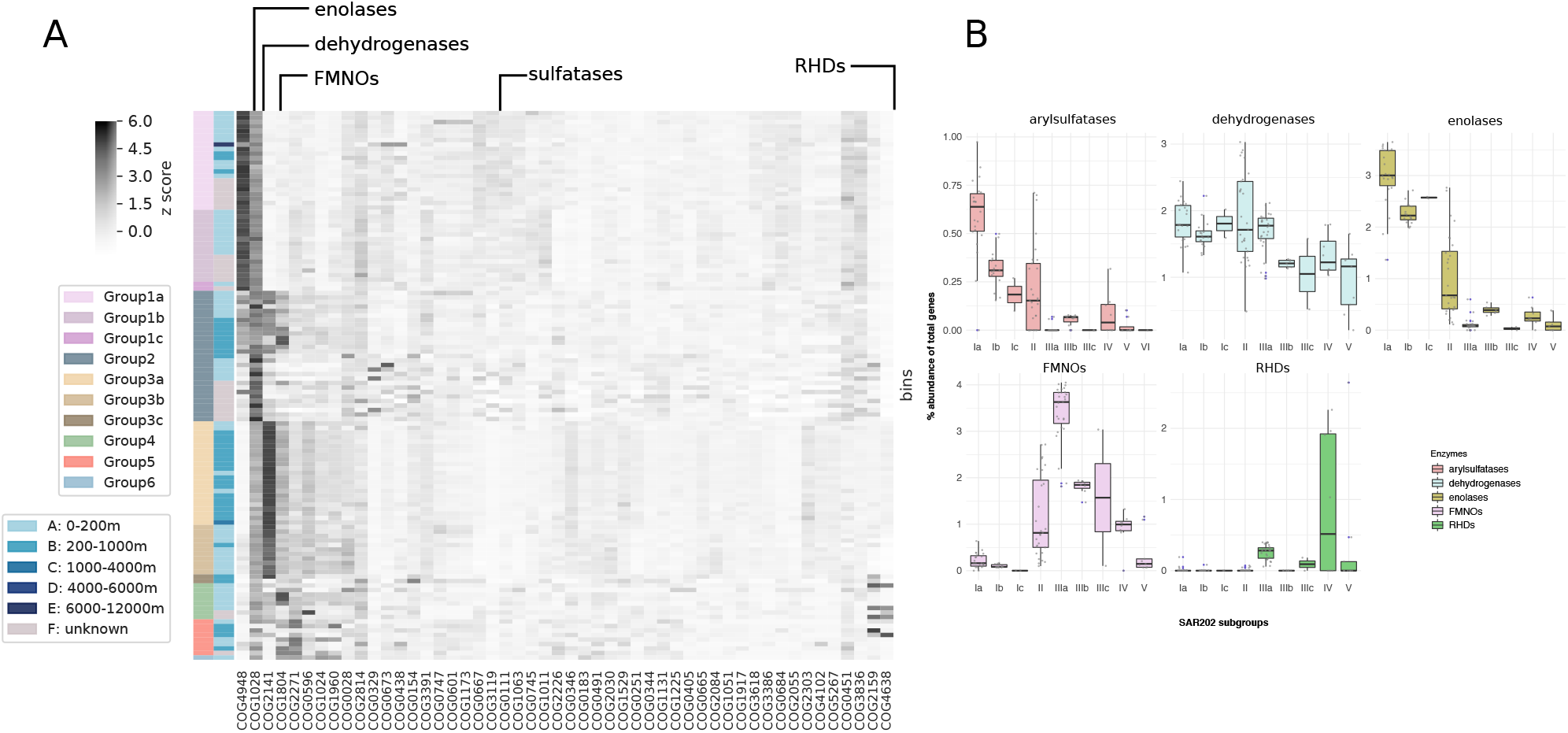
**(A)** Heatmap of most abundant COG categories in SAR202 genomes categorized by sub-groups. The first column of color bars indicates different SAR202 subgroups and the second column of color bars indicate the depth of samples from which the SAGs or the MAGs were obtained. The number on the heatmap color gradient indicates z scores of percent abundance of total number of genes. **(B)** Distribution of the major paralog expansions among the SAR202 subgroups.

A correlation matrix of the top 50 most abundant COG categories showed that the expansions of the five major paralog families discussed above are linked to broad shifts in metabolism (Fig. 3). For example, COG3391, COG4102, and COG5267 are all uncharacterized conserved proteins. COG0747, COG0601, and COG1173 are components involved in dipeptide transport. We interpret these patterns as evidence that the ancient paralog expansions described above accompanied metabolic reorganization and specialization in the SAR202 subclades.

**Figure 3:**
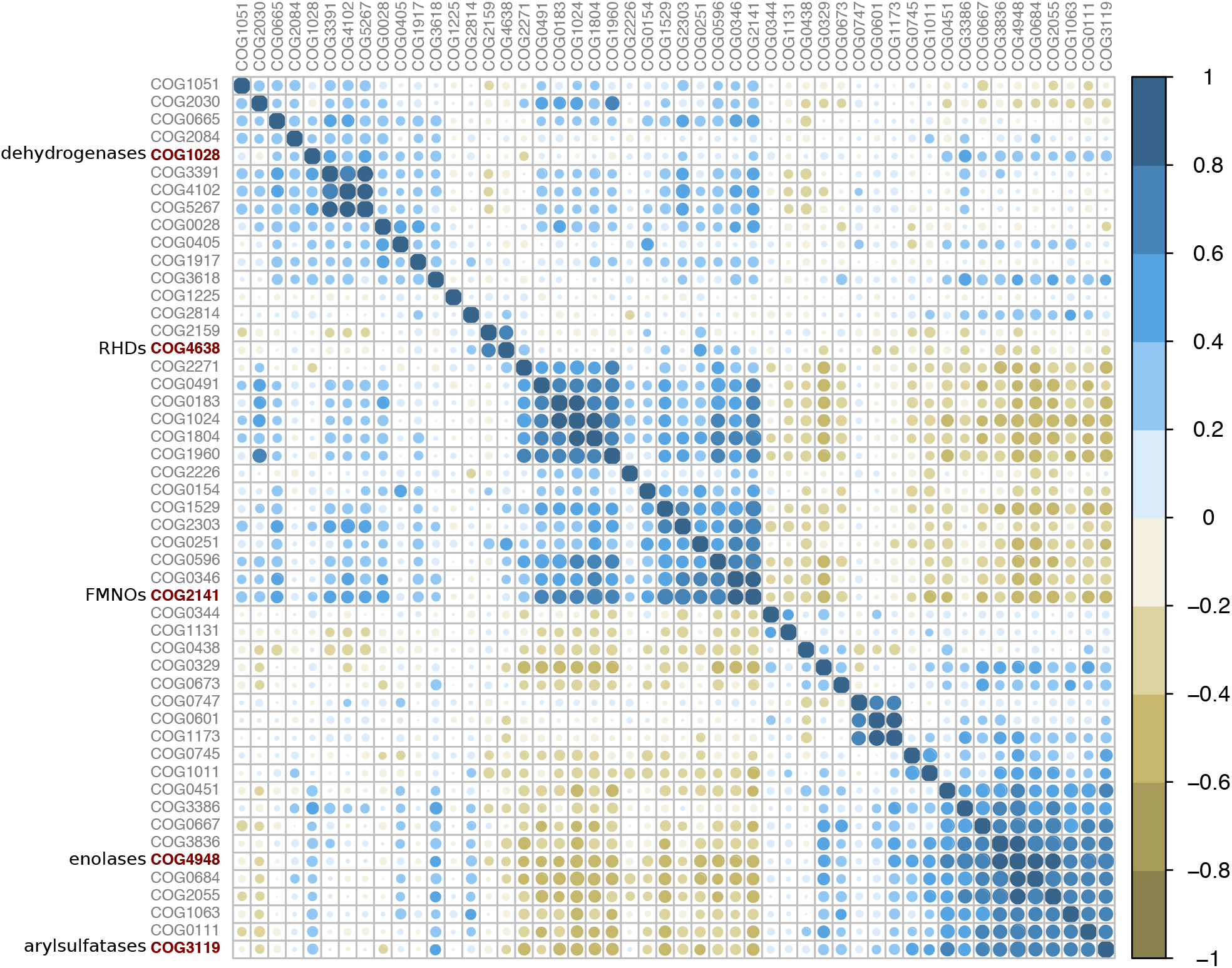
Correlations among top 50 most abundant COG functional categories, demonstrating that the major paralog expansions identified in Figure 2 are linked to other expanded families of proteins, indicating metabolic specialization.

### The diversification of flavin-dependent monooxygenases in Group III

An expansion and radiation of diverse FMNO members in Group III SAR202 was previously reported (10). We found further support for this conclusion in this broader analysis of SAR202 diversity, and also observed elevated numbers of FMNO paralogs in Groups II and IV. The number of paralogous FMNO copies ranged from 1 and 114, with members of Group IIIa encoding the highest numbers and the greatest relative abundances, up to 4% when normalized to total number of resolved genes (Fig. 2B). FMNOs were also present in other SAR202 subgroups, at lower copy numbers. Group 1 encode the fewest copies of FMNOs; in some genomes this number approaches zero. The five most abundant FMNOs were annotated as: alkanal mono-oxygenase alpha chain (23% of all annotations); limonene 1,2-monooxygenase (21%); phthiodiolone/ phenolphthiodiolone dimycocerosates ketoreductase (13.9%); F420-dependent glucose-6-phosphate dehydrogenase (13.7%); and alkanesulfonate monooxygenase (7.2%).

Because automatic annotation can sometimes fail to assign proper function to the genes, we built a maximum likelihood (ML) phylogenetic tree of all extant FMNOs identified in databases to better visualize the functional diversity of the FMNOs (Fig. 4A). We identified five broadly-classified functional groups: F420-dependent tetrahydromethanopterin reductases, alkanal monooxygenases, nitrilotriacetate monooxygenases, alkanesulfonate monooxygenases, and pyrimidine monooxygenases (RutA). Most fall into the alkanal and F420-dependent monooxygenases. The SAR202 F420-dependent monooxygenases are highly diverse and appear to be paraphyletic. It remains to be determined whether SAR202 can synthesize coenzyme F420.

**Figure 4:**
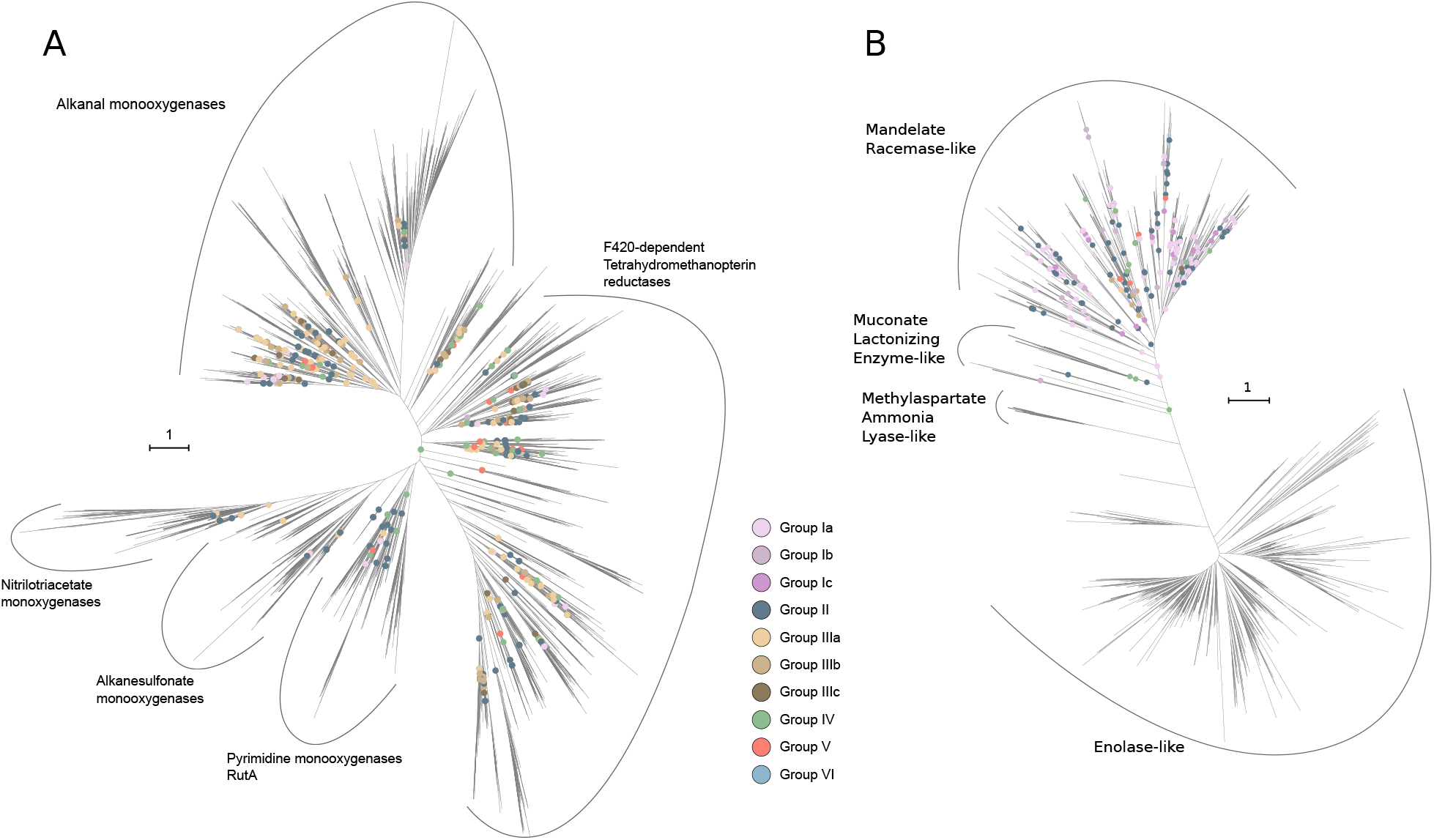
**(A)** Phylogenetic tree of the FMNO superfamily of enzymes. Internal nodes marked with colored circles indicate points of attachment for SAR202 lineages. The deep positions of the SAR202 nodes suggest that a substantial part of enzyme diversity in the FMNO superfamily is found in SAR202. The cluster of Group IIIA nodes deep in the alkanal monooxygenase subclade suggest that these enzymes, in particular, may have evolved in SAR202. **(B)** Phylogenetic tree of the enolase superfamily of enzymes. SAR202 paralogs branch deeply and are confined to the madelate racemase-like enzyme sub-family of enolases. Scale bar represents the number of amino acid substitutions.

Type II Baeyer-Villiger monooxygenases were found in Group IIIa SAR202 as described previously (10) and fall into the broad category of alkanal monooxygenases. The alkanal monooxygenases formed a monophyletic clade with deepest nodes belonging to Group IIIa genes (Fig. 4A). This pattern indicates that this sub-family of enzymes may have originated within SAR202 Group IIIa.

### The Group I & II enolase paralog expansion, an adaptation to unlock chiral diversity in DOM resources?

We observed an expansion of diverse enolase superfamily paralogs in Groups I and II (Fig. 2A, 2B, and 4B). The presence of enolase paralogs in SAR202 genomes was first noted in MAGs obtained from a northern Gulf of Mexico ‘dead zone’ (14). Annotations of five most abundant SAR202 enolases are: D-galactonate dehydratase (52.9% of all annotations); L-rhamnonate dehydratase (16.4%); starvation-sensing protein RspA (10%); mandelate racemase (6.8%); and L-Ala-D/L-Glu epimerase (5.4%).

The numbers of enolase paralogs in Group 1 ranged from 4 to 75 (1.3 to 3.5% of total genes found in each subclade); other SAR202 clades appear to encode very few copies of this enzyme (Fig. 2B), with the exception of Group II SAR202, which encode both FMNO and enolase paralogs, in roughly equal abundances (Fig. 2B). Enzymes of the enolase superfamily catalyze mechanistically diverse reactions such as racemizations, epimerizations, *β*-eliminations of hydroxyl or amino groups, and cycloisomerizations, but all the known reactions they catalyze involve abstraction of an *α*-proton from carbons adjacent to carboxylic acid groups and stabilization of the enolate anion intermediate through a divalent metal ion, usually Mg^2+^ (24, 25).

Muconate cycloisomerases were also detected in SAR202, although they constitute a small fraction of the enolases found. They belong to the muconate lactonizing enzyme (MLE) family and are involved in breaking down of lignin-derived aromatic compounds, catechols, and protocatechuate to produce intermediates that are used in the citric acid cycle (26, 27). It is worth noting that, although Group I members predominantly encode a large diversity of enolase family enzymes, some Group III members also encode a few of these genes, the majority of which are mandelate racemases (Fig. 2B and 4B).

A phylogenetic tree was constructed to highlight the diversity and functions of enolase family enzymes found in Group I SAR202 genomes. Enzymes within this superfamily can be divided into four categories: enolases, mandelate racemases, muconate lactonizing enzymes, and methylaspartate ammonia lyases (Fig. 4B). Nearly all of the enolases in SAR202 belong to the mandelate racemase family. Enzymes within this family include mandelate racemase, galactonate dehydratase, glucarate dehydratase, idarate dehydratase and similar enzymes that can either interconvert two stereoisomers or perform dehydration reactions (24).

Enzymes that can interconvert between *R* and *S* forms (stereoisomers) could vastly improve the fitness of an organism by making it able to utilize both compounds. For example, organisms that encode mandelate racemase (MR) in their genomes can interconvert between *(R)*-mandelate and *(S)*-mandelate, the latter of which is the first compound in the mandelate and hydroxy-mandelate degradation pathways (28). We postulate the expansion of diverse enolase superfamily paralogs in Groups I and II is an adaptation to metabolize organic compounds that are recalcitrant to oxidation because of chiral complexity. In the discussion section, we further explore the ramifications of these observations.

### Sulfatases in Group I and II members

Sulfatases in SAR202 were first reported in a study on dead zones in Gulf of Mexico (14). We also detected a large number of genes belonging to COG3119 (AslA, Arylsulfatase A) and related enzymes classified in inorganic ion transport and metabolism predominantly in Group I and II bins (Fig. 2B). Arylsulfatases and choline sulfatases can hydrolyze sulfated polysaccharides such as fucoidan produced by marine eukaryotes (algae or fungi). These enzymes are expressed intracellularly by a species of marine fungus (29), and are also found in marine *Rhodobacteraceae* that are mutualists of marine eukaryotes (30). Marine brown algae, such as *Macrocystis*, are known to produce fucoidans, which consist of *α*-L-fucosyl monomers (31). We speculate that SAR202 Groups 1 and 2 could be utilizing arylsulfatases to break down similar sulfated polysaccharides produced by the algae in the upper water column.

### Ring-hydroxylating dioxygenases in Group IV, a molecular arsenal to break down aromatic compounds

One of the enzyme families that seems to be disproportionately expanded in SAR202 belongs to COG4638, annotated as “phenylpropionate dioxygenases or related ring-hydroxylating dioxygenases, large terminal subunit”. Enzymes belonging to the ring-hydroxylating dioxygenases (RHDs) family occur as monomers of subunits alpha and beta (*α*_2_*β*_2_ or *α*_3_*β*_3_) (32). The *α* subunit of RHDs contains a Rieske [2Fe-2S] center that transfer electrons to iron at the active site while the *β* subunit is thought to play a structural role in the enzyme complex (32). Members of SAR202 Group IV harbor a large number of these RHDs, ranging from 1 to 62 paralogous copies for subunit *α* (COG4638) and 1 to 3 for subunit *β* (COG5517). Given that there are more *α* than *β* subunits, it appears that most of the RHDs in Group IV function as monomeric RHDs.

Of the 365 RHD *α* subunits found in SAR202, 136 copies came from Group 4. OSU_TB11, a Group 4 SAR202, encodes the highest relative abundance of RHDs at 50 (2.64%) of all genes in its genome (Fig. 2B). A sponge symbiont member of Group IV (MPMJ01) (22) encodes the largest number of copies of RHDs (62 copies and 1.96% of its genes), but it also has one of the largest genomes, 3.22 Mbp. Most of the RHDs were annotated as: phthalate 4,5-dioxygenase oxygenase subunit (38.9%), phenoxybenzoate dioxygenase subunit alpha (26%), 3-phenylpropionate/cinnamic acid dioxygenase subunit alpha (20.5%), or carbazole 1,9a-dioxygenase, terminal oxygenase component (8.2%).

While the vast majority of the RHDs are annotated as “phthalate 4,5-dioxygenases”, it is unlikely that phthalates are common substrates in the ocean. Most of Group IV SAGs and MAGs were recovered from euphotic zone samples; all bins originated from ≤ 200 m depth. We speculate these enzymes are used to metabolize other mono- or polycyclic aromatic compounds that are mainly released by phytoplankton, providing Group IV SAR202 with energy and carbon.

A recent paper showed that some of the SAR202 members encode large numbers of RHDs in their genomes, which were likely acquired by horizontal gene transfer (HGT), and speculated they play a role in the catabolism of resistant DOM of terrestrial origin (16). We found Group IV MAGs containing copies of RHDs predominantly in samples from coastal regions of the Indian Ocean and Red Sea, and the Southern Ocean, near Antarctica (Fig. S1).

### Rhodopsins in epipelagic Group I and II SAR202

Twenty-eight genomes, all from samples obtained from water depths shallower than 150 m, encoded proteorhodopsins, one of which was a heliorhodopsin. Most of the type-1 rhodopsins were found in members of Group Ia, Ib, Ic, and Group II, which we report are prevalent in the euphotic zone. The single heliorhodopsin, which was found in a Group II genome, is related to a recently described group of heliorhodopsins (35). Using the backbone tree from that study (35), the SAR202 Type-1 rhodopsins were placed close to previously known proteorhodopsins and the sole heliorhodopsin was placed deep within the newly described heliorhodopsins (Fig. S2 and S3).

### Depth stratification and biogeography indicate niche specialization is correlated with expansions of paralogous gene superfamilies in SAR202

Group I genomes, including those that encoded rhodopsins, were mostly isolated from the epipelagic (0-200 m), whereas the Group III members were mainly retrieved from the mesopelagic (200-1000 m) (Fig. 2). We further analyzed a variety of data types and found that the major SAR202 Groups have different depth ranges (Fig. 5). The oceanic water column vertical gradients of light (PAR), inorganic nutrients and organic matter quality and quantity establish specialized nutritional niches. The vertical stratification of SAR202 groups with the evidence described above for metabolic specialization, suggests that SAR202 diversified to specialize in resources that vary across the water column.

**Figure 5:**
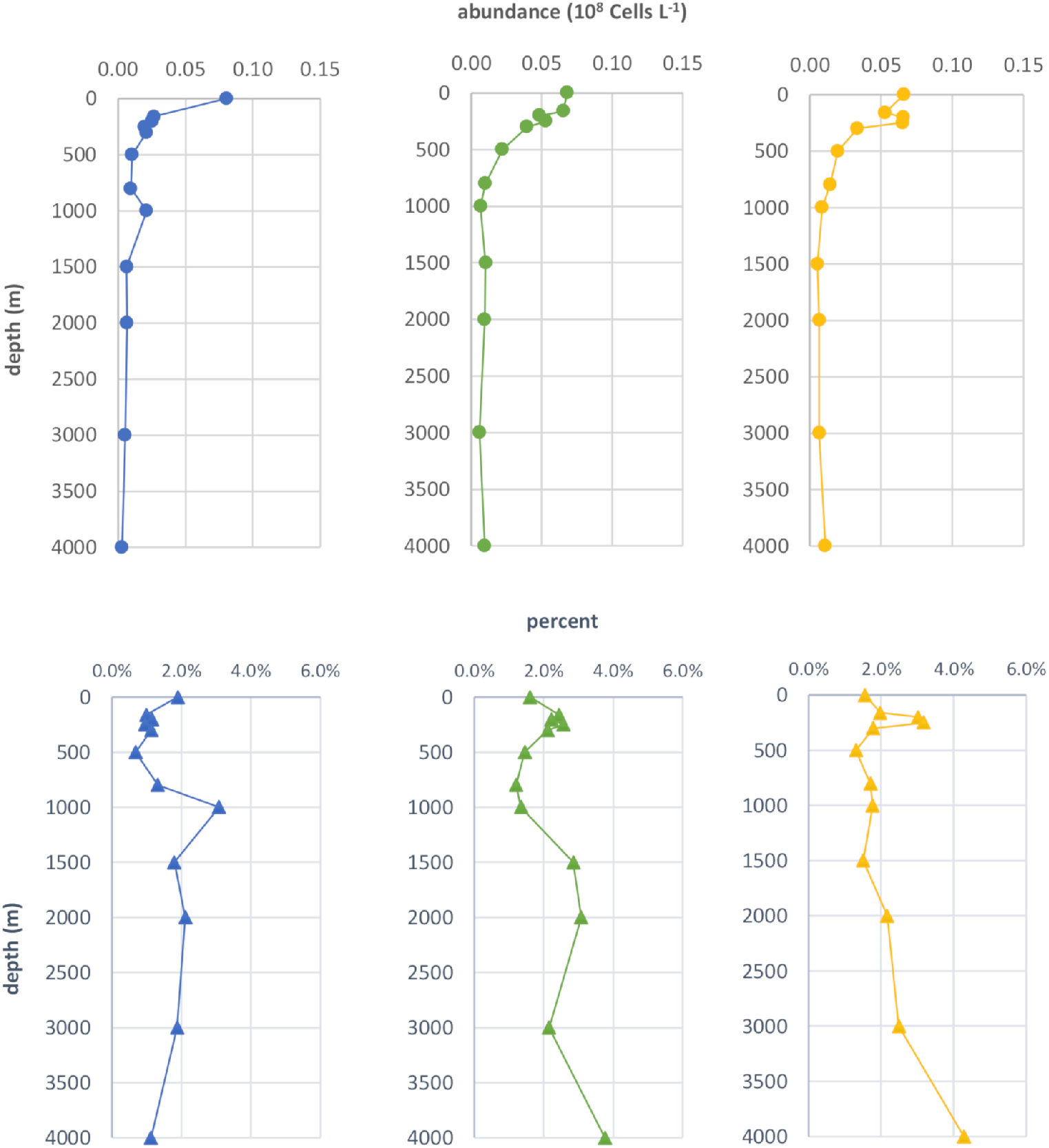
Depth profiles showing SAR202 Group I abundance (blue circle and line); SAR202 Group II abundance (green circle and line) and SAR202 Group III abundance (yellow circle and line) as determined by FISH group-specific oligonucleotide probes. Depth profiles showing SAR202 Group I percent contribution to total bacterioplankton determined by DAPI cell counts (blue triangle and line); SAR202 Group II percent contribution to total bacterioplankton (green triangle and line) and SAR202 Group III percent contribution to total bacterioplankton (yellow triangle and line).

### Fragment recruitment analyses

Metagenome fragment recruitment showed that Group I members are most abundant in the epipelagic (from surface to 200 m); Group III recruited more reads from meso, bathy, abysso and hadalpelagic samples, and Group II recruited reads from the surface through the mesopelagic (Fig. 6, S4, and S5). In TARA Oceans metagenomes, Group I members, most notably Ib, were relatively more abundant in the epipelagic (5-80 m in the Indian Ocean, 5-60 m in the Mediterranean Sea, 100-150 m in the South Atlantic Ocean, and 115-188 m in the South Pacific Ocean) (Fig. S4). However, despite decreasing with depth, their abundance didn’t reach zero, indicating populations persist in the deep ocean. In waters overlying the Japan and Mariana Trenches, Group I members (particularly Ib), were abundant only near the surface.

**Figure 6:**
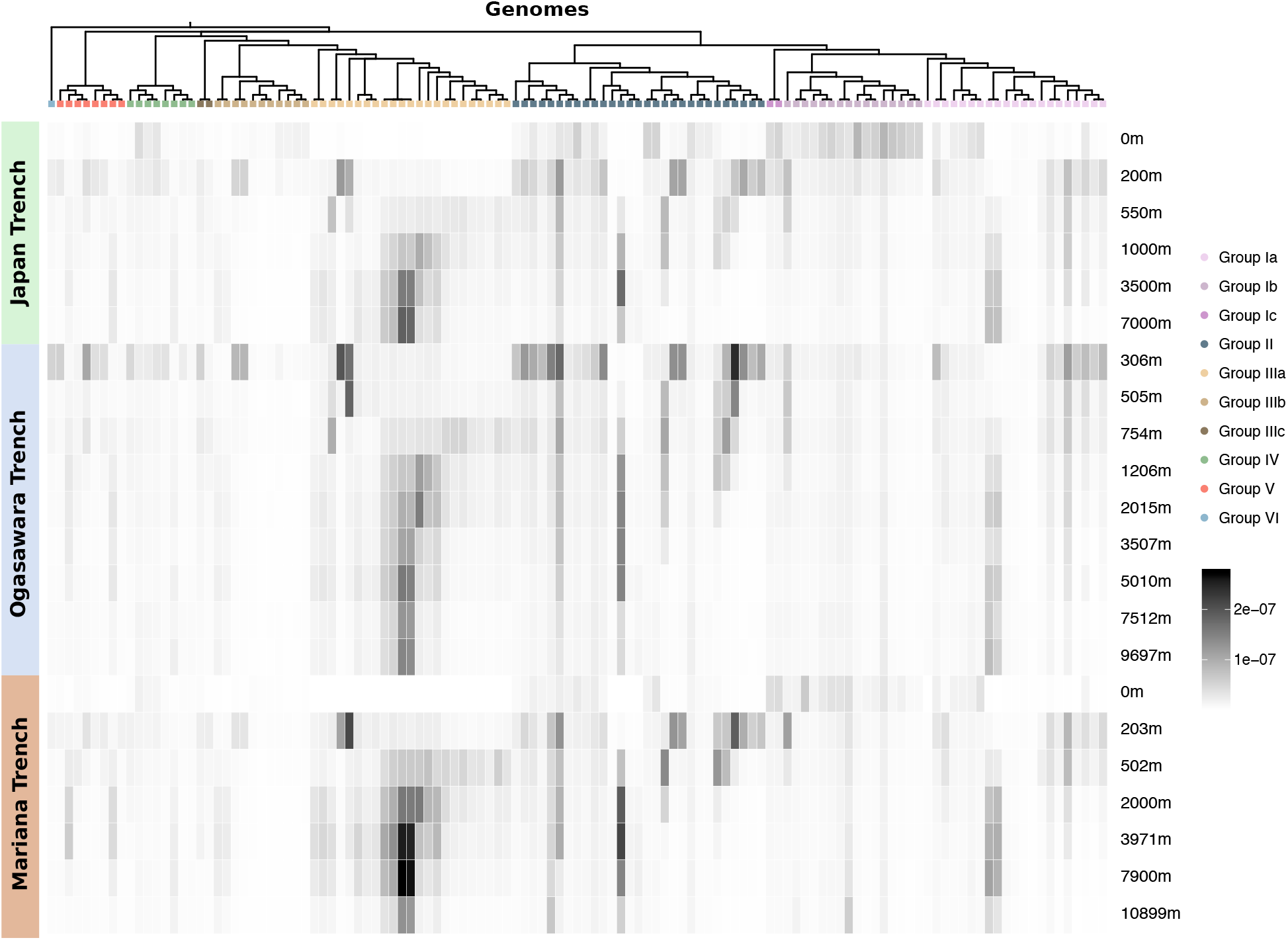
Fragment recruitment analysis of metagenomic reads from three deep-ocean trenches against the SAR202 genomes. Arrangement of SAR202 genomes follows the branching order in the Bayesian phylogenomic tree shown in Figure 1. Recruitment is calculated as the number of bases of metagenomic reads aligned against SAGs or MAGs normalized by total number of bases present in a given metagenomic sample. The intensity of shading represents the degree of recruitment.

There is a noticeable absence of Group IIIa members in upper water column above 200 m in the Northwestern trenches metagenomes (Fig. 6), above 250 m in the TARA Oceans metagenomes (Fig. S4), and above 200 m in BATS metagenomes (Fig. S5). They are most abundant in deeper layers (600-1000 m in the Indian Ocean, 590-800 m in the North Atlantic Ocean, 700-800 m in the South Atlantic Ocean, 375-650 m in the North Pacific Ocean, 350-696 m in the South Pacific Ocean, and 790 m in the Southern Ocean) (Fig. S4). Group IIIa members are found almost exclusively below 200 m (200-7000 m at Japan Trench, 306-9697 m at Ogasawara Trench, and 203-10899 m at Mariana Trench). Members of Group IIIb, however appear to be more abundant in the upper water columns and less so in the deeper zones in two metagenome datasets (Fig. 6 and S4).

Group II members seem to occupy transitional zones between those occupied by Group I and Group III members (for example, 270-600 m in the Indian Ocean, 250m in the North Atlantic Ocean, and 40-450 m in the North Pacific Ocean). However, the zones occupied by Group II members seem to largely overlap with those of both Group I and Group III members as well (Fig. 6 and S4). Group II members are again found to occupy intermediate depths in the Northwestern Pacific Ocean trenches (200-1000 m at Japan Trench, 306-1206 at Ogasawara Trench, and 203-502 m at Mariana Trench). Some Group II members are found in wider depth ranges, with one found to be quite abundant in deepest water samples in all three trenches (Fig. 6).

### Group I, II and III Florescence in Situ Hybridization Profiles

The first group-specific oligonucleotide probes for SAR202 Groups I, II and III were developed and used to count cells throughout the BATS water column to 4000 m in July 2017 (Fig. 5). All three groups were detected in significant numbers throughout the water column, summing to about 5% of total bacteria near the surface and up to 10% at 4000 m. Group I SAR202 cell numbers peaked in the epipelagic and dropped off sharply below the euphotic zone (100 m), whereas both Group II and III had a broader distribution across the epipelagic, peaking sharply within the upper mesopelagic zone at ~ 250 m, as reported previously. When plotted as relative abundance (lower panels, Fig. 5), the direct cell count data was consistent with the observations from metagenome recruitment, which are also presented in relative units.

### SAR202 FMNO gene relative abundance is correlated with depth

The relative abundance of all TARA FMNO genes (Fig. S8C), and SAR202 specific FMNOs, was correlated with depth (Fig. 7C), with Pearson r values for the latter of 0.87 (P=9.6e^−75^). From these results, it was clear that FMNOs appear to be more functionally important in the deeper oceans.

**Figure 7:**
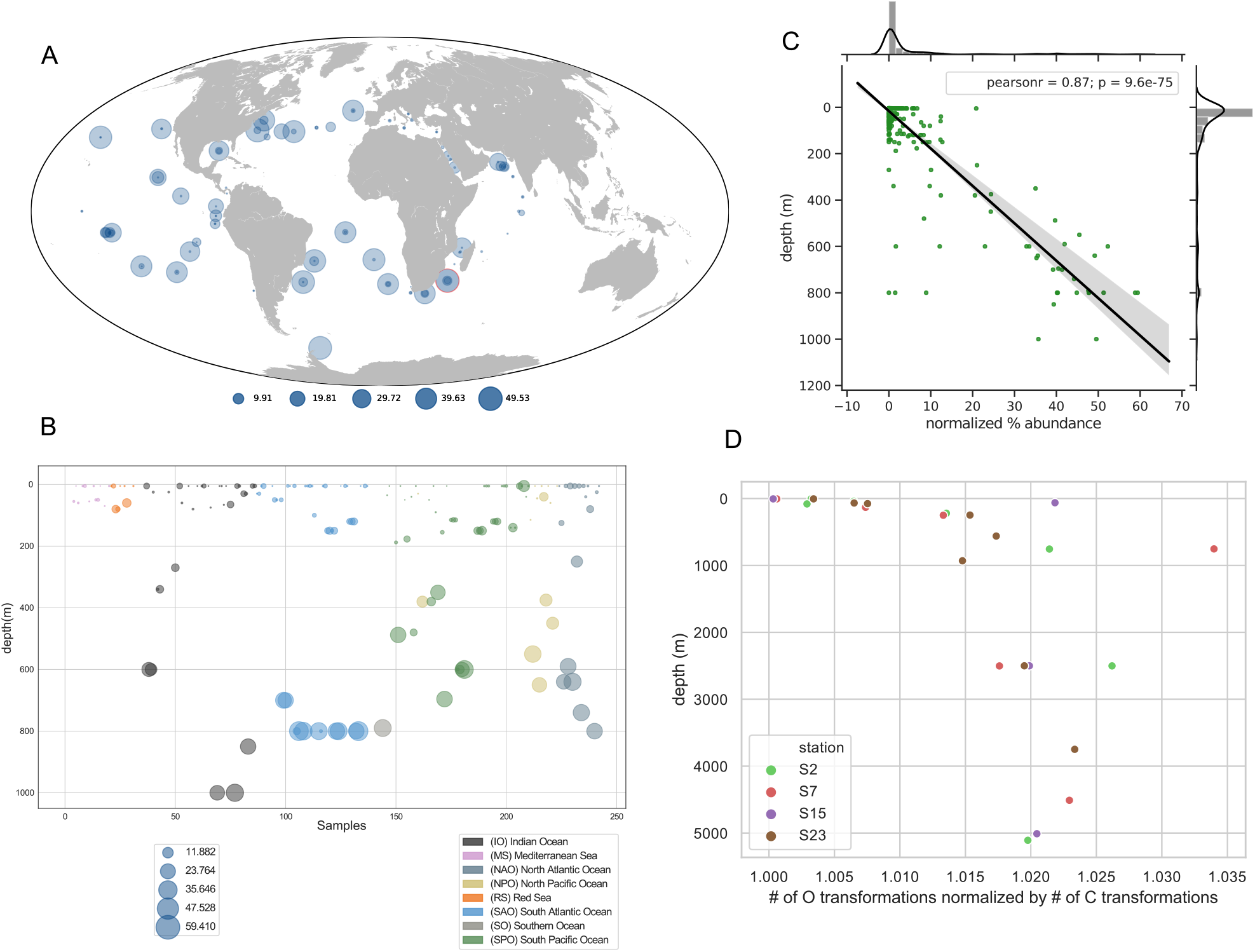
**(A)** World Map showing relative abundances of SAR202-specific FMNOs in TARA Oceans metagenomes. Sample with highest relative abundance is highlighted in red circle. **(B)** SAR202-specific FMNOs relative abundances vs. depth in TARA oceans metagenomes. **(C)** Normalized FMNO abundances in SAR202 are highly correlated with depth in TARA Oceans metagenomes. Normalization of FMNO abundances was obtained by dividing total SAR202 FMNOs by total SAR202 single-copy genes found in each sample. **(D)** The ratio of observations of organic metabolites with mass : charge ratio (m/z) that differ in mass by one oxygen, to observations that differ in mass by one carbon, in FTICR-MS data from deep ocean marine DOM samples collected from the Western Atlantic. The stations ranged from 38° S (station 2) to 10° N (station 23). Across the full dataset, the most common m/z difference observed corresponds to one carbon atom of mass. The data show that transformations corresponding to the addition of a single oxygen atom, as would be catalyzed by a flavin-dependent monooxygenase, become relatively more frequent in the dark ocean. Of several patterns predicted from a previous study (10), this one alone showed a consistent trend.

Because it appeared that FMNOs are abundant in SAR202 members originating from the bathy- and abysso-pelagic, we checked to see if the relative abundances of FMNOs in SAR202 genomes correlated with depth. Fig. S6A shows a significant positive correlation between FMNO relative abundance vs. depth and Fig. S6B shows weak but significant negative correlation between enolase abundances vs. depth. These data indicate that FMNOs are mostly abundant SAR202 cells from deep waters, whereas the enolases are more abundant in shallow water ecotypes.

The analysis in Fig. 7D tests the prediction that molecules differing by the addition of a single oxygen atom, as expected from the chemical mechanism of FMNO enzymes, should be more abundant in the deep ocean. In the plot, the ratio between the number of m/z observations that differ in mass by one oxygen, to observations that differ in mass by one carbon, increases dramatically below the epipelagic. In the model we presented previously, cells are presumed to enzymatically modify resistant DOM compounds, channeling some to catabolism, while exporting from the cell molecules that cannot be further degraded (10).

### Enolase abundances show weak correlation with depth

Because enolases appear to be a notable feature of SAR202 SAGs and MAGs from the upper water column, we assessed whether relative enolase abundances were also correlated with depth. Fig. S6B shows that there is a slight negative correlation between the % abundance of enolase genes in MAGS and SAGS and the depth they were recovered from, but SAR202 enolases in the TARA Oceans metagenomic data show a somewhat positive correlation with depth (Pearson r value of 0.6, P=1.4e^−25^) (Fig. S7). This was surprising because we reasoned that the enolases might be involved in breaking down more labile compounds found in the upper water column based on the genomic data and expected higher abundances of enolases in the samples from upper water columns. One reason for this discrepancy could be biased sampling of MAGs from TARA Oceans metagenome samples. We selected 43 TARA samples to re-assemble based on SAR202 abundances; some samples from deeper regions that we did not assemble could harbor uncharacterized SAR202 subgroups that encode a large number of enolases.

## Discussion

Pangenome analysis confirmed earlier reports and uncovered further evidence of ancient expansions of paralogous enzymes in the SAR202 clade (Fig. 2B, 4A, 4B). The paralogous gene families were correlated with deep branches in the SAR202 genome tree, which divide the clade into six subgroups. Metagenome analyses, and cell counts made with FISH probes, showed that several of the SAR202 groups are vertically stratified through the water column, suggesting niche specialization (Fig. 6). Collectively, these patterns amount to strong evidence that the early evolutionary radiation of SAR202 into subgroups was accompanied by metabolic specialization and expansion into different ocean niches.

It is striking that the major paralog expansions in SAR202 suggest three different metabolic strategies, each potentially targeting a different class of semi-labile DOM compounds. In the hypothetical schemes we developed, the evolutionary diversification of paralogous enzyme families was driven by selection favoring substrate range expansion. We found support for this scheme in evidence these gene lineages arose early in evolution. While deep internal nodes for these genes in tree topologies could result from the recruitment of paralogs by horizontal gene transfer, the rarity of near gene neighbors across the tree-of-life favors the explanation that most of the paralog diversity arose within SAR202 by gene duplication during evolution. If this interpretation is correct, it implies that much of the functional diversity in two major enzyme families, the alkanal monooxygenases within the FMNO superfamily and madelate racemases within the racemase superfamily, may have originated within SAR202. This is apparently not the case for the Group IV dioxygenases, for which there is evidence of acquisition by HGT (16).

Surprisingly, because SAR202 have the reputation of being deep ocean microbes, the ecological data we gathered revealed that Group I SAR202 are mainly epipelagic, and harbor large and diverse families of enolase paralogs. We interpret this proliferation of enolase superfamily paralogs as evidence that these organisms have evolved to metabolize organic matter that is resistant to oxidation because of chiral complexity. Enolase superfamily enzymes remove the α-proton from carboxylic acids to form enolic intermediates, which can rotate on the axis of the double bond of the intermediate, with stereochemical consequences (24). These enzymes catalyze racemizations, β-eliminations of water, β-eliminations of ammonia, and cycloisomerizations. Chemical oceanographers have recognized a role for molecular chirality in diagenesis, reporting that the ratio of D- to L-aspartic acid uptake by prokaryotic plankton increases by two to three orders magnitude between surface and deep mesopelagic waters in the North Atlantic (36). This has been interpreted as evidence that mesopelagic prokaryotic plankton are using bacterial cell wall–derived organic matter because the bacterial peptidoglycan layer is the only major biotic source of significant of D–amino acids in the ocean (37). However, information about D-amino utilization by marine microbes remains limited (38).

The possibility that SAR202 harness paralogous enzymes of the enolase superfamily to metabolize compounds that are resistant because of chirality is a powerful concept. We propose that chiral complexity defines a class of resistant compounds, and that enolases are an innovation that makes this DOM accessible to degradation by reducing the number of enzymes needed to degrade it. The number of enantiomers of a compound increases by 2^n^, where n is the number of chiral centers. Thus, a single compound with three chiral centers might in principle require eight enzymes to recognize all stereoisomers. However, if the three chiral centers were racemized by enolases, then only four enzymes would be required – one degradative enzyme and one enzyme to racemize each of the chiral centers. Spontaneous racemization might play a role in increasing the chiral complexity of DOM and thereby transitioning it to more resistant forms, but it might also originate in biological complexity, much of which is unexplored. The role for enolases that we propose evokes the *molecular diversity hypothesis* by speculating there is a relationship between the complexity of DOM and its resistance to degradation. Most often, the *molecular diversity hypothesis* is used to explain the relationship between the dilution of DOM and its susceptibility to degradation.

We speculate that Group I SAR202 are specialized to harvest a fraction of DOM molecules that are semi-labile because of unusual chiral structures. Group II SAR202, which are most abundant in the mesopelagic, maintain both the enolase and FMNO enzyme families in equal abundances, suggesting they use both DOM resources – chirally complex organic matter and compounds that can be catabolized via monooxygenases - in this intermediate water column zone. Earlier studies have demonstrated that, in addition to a DOC concentration decreasing with ocean depth, the abundance of diagenetically altered DOM compounds increases below the euphotic zone (39–41). In bathypelagic, abyssopelagic and hadalpelagic regions, Group III dominate, presumably indicating that molecules susceptible to oxidation by FMNOs become one of the few remaining harvestable DOM resources at these depths. In this scenario, SAR202 diversified strategically to exploit multiple different classes of resistant carbon compounds in niches distributed throughout the water column. The positions and separation of the subclades in trees, and the diversity of the enzymes involved, suggest this evolution occurred early in SAR202 history. Close examination of Fig. 6 shows that there are more finely structured patterns of congruence between tree topologies and depth range than the broad patterns we focus our discussion on. For example, some lineages of Group Ia were consistently observed in bathypelagic, and some Group II near the surface. It is apparent that more complex relationships between ecology, evolution and metabolism remain to be explored in SAR202.

This study confirmed previous reports of expansions of FMNO enzymes in Group III genomes recovered from the deepest ocean regions (10), and RHD enzymes in Group IV genomes from coastal sites. Both FMNO and RHD enzymes are powerful oxidases implicated in the catabolism of resistant compounds such as sterols and lignins. The expansion of these enzyme families is proposed to have enabled SAR202 to exploit new niches defined by these DOM resources. In the case of Group IV this would be lignins and other aromatic compounds of terrestrial origin, whereas Group III is proposed to partially oxidize a wide variety of recalcitrant molecules, including perhaps sulfonates and heterocyclic compounds. It has been hypothesized that the partial oxidation of these compounds might produce more recalcitrant compounds that accumulate RDOM.

The genome-enabled hypotheses we propose will be challenging to test, but nonetheless should be studied because the organic carbon pool in question is so large. Deep-ocean regions beyond the reach of sunlight contain an estimated 662 Pg of DOC (1), which ranges in quality between LDOM and RDOM (3, 42). If our hypotheses are correct, this pool would be much larger if cells had not evolved strategies to oxidize many forms of resistant DOM. In principle, the modern RDOM pool would become much smaller if contemporary cells evolved mechanisms to oxidize it, with catastrophic consequences for the environment.

The complexity of DOM presents many challenges to proving these hypotheses. Thus far, DOM chemical structures have not been resolved with sufficient accuracy to support a detailed accounting of compounds and corresponding pathways of microbial catabolism. An example of these problems is the issue of chemical enantiomers, which have identical empirical formulas, making them perhaps the most difficult challenge. In brainstorming these challenges, we encountered one success (Fig. 7D) which illustrates both the difficulty of the task and the hope for finding solutions. Future work might focus both on the composition of DOM and the activities of cells that are not yet cultured in laboratories.

## Materials and Methods

Methods for metagenomic library preparation and sequencing, single-gene phylogenetic and phylogenomic analyses, direct cell counts and fluorescent in-situ hybridization of SAR202 can be found in the supplemental online document.

### Sample collection and sequencing of single amplified genomes and shotgun metagenomic sequencing from the three trench sites

SAG generation was performed using fluorescence-activated cell sorting and multiple displacement amplification at Bigelow Laboratory Single Cell Genomics Center (SCGC; scgc.bigelow.org), as previously described (43). Selection for genomic sequencing was aimed at representing the diverse SAR202 subgroups based on their 16S rRNA phylogenetic tree placement and 10 single-cell amplified genomes (SAGs) were selected for genomic sequencing based on the phylogenetic placement (data not shown). They originate from samples from three deep-sea trenches in the Northwestern Pacific Ocean: Mariana, Japan, and Ogasawara Trenches. Water samples from the central part of the Izu-Ogasawara (Izu-Bonin) Trench (29°9.00’ N, 142°48.07’ E, 9776 m below sea surface [mbs]) were obtained using Niskin-X bottles (5-liter type, General Oceanics) during a total of two dives of the *ROV ABISMO* during the Japan Agency for Marine-Earth Science & Technology (JAMSTEC) *R/V Kairei* KR11-11 cruise (Dec 2011). Water samples from the southern part of the Japan Trench (36° 5.88’ N, 142° 45.91’ E, 8012 mbs) was obtained by vertical hydrocasts of the CTD-CMS (Conductivity Temperature Depth profiler with Carousel Multiple Sampling system) with Niskin-X bottles (12-liter type, General Oceanics) during the JAMSTEC *R/V Kairei* KR12-19 cruise (Dec 2012). From the Challenger Deep of the Mariana Trench Water samples except for the trench bottom water were taken by Niskin-X bottles (5-liter type) on the *ROV ABISMO* and the trench bottom water was obtained by a lander system (44) during the JAMSTEC *R/V Kairei* KR14-01 cruise (Jan 2014). Samples for SAG generation were stored at −80°C with 5 % glycerol and 1 x Tris-EDTA buffer (final concentrations) (45). For the shotgun metagenomic library construction, Microbial cells in approximately 3-4 L of seawater were filtered using a cellulose acetate membrane filter (pore size of 0.22 μm, diameter of 47 mm) (Advantec, Tokyo, Japan).

Four SAGs were sequenced at SCGC and six SAGs were sequenced at Center for Genome Research and Biocomputing (CGRB) at Oregon State University after NexteraXT sequencing libraries were prepared at JAMSTEC. Sequencing libraries for SAGs obtained from the Mariana Trench site was directly synthesized with Nextera XT DNA Library Preparation Kit (Nextera XT) as described previously (46). The amplification cycle for the construction of these libraries was 17 except the case of AD AD-812-D07 with 12 cycles of amplification.

### Genome assemblies, binning, and annotation

Illumina library preparation, sequencing, de novo assembly and QC of SAGs AC-409-J13, AC-647-N09, AC-647-P02 and AD-493-K16 were performed by SCGC, as previously described (43). For the remaining six SAGs, raw sequences were first quality trimmed using Trimmomatic tool (47). Four SAGs were assembled individually using SPAdes assembler version 3.9.0 (48) with “–careful and -sc” flags. Due to cross-contamination present in a second batch of 6 SAGs sequenced, they were co-assembled using metaSPAdes, then CONCOCT was used to separate the contigs from each SAG into respective bins. CheckM analysis of the bins showed that contamination levels in each identified bin were very low (below 0.2%) and the 6 SAGs are from very divergent clades, so that they can be easily separated by differential coverage binning approach.

Raw sequences from 17 metagenomics samples from Bermuda Atlantic Time-series Study (BATS) and 43 metagenomic samples from TARA Oceans expedition were quality trimmed using Trimmomatic and individually assembled using metaSPAdes version 3.9.0 (49). The 43 TARA Oceans metagenomes chosen contain at least 1% of relative SAR202 abundance based on metagenomics tag (miTAG) sequence data (50) (Supplemental Table 2).

All metagenomics contigs larger than 1.5 kbp were separated using metabat (51) to gather potential SAR202 bins. Metabat requires the use of multiple samples to calculate contig abundance profile in the samples. For TARA Oceans metagenomes, in order to generate abundance profiles, contigs were mapped against a minimum of 10 TARA oceans metagenome samples chosen randomly (including the sample from which the contigs were assembled) using BBmap (http://sourceforge.net/projects/bbmap/). For BATS metagenomes, BBmap was also used against all 17 metagenomes to generate config abundance profiles. Identities of the resulting bins were checked for presence of 16S rRNA gene sequence matching known SAR202 sequences from Silva database release 128. In cases where there were no 16S rRNA genes in the bins, concatenated ribosomal protein phylogenies were constructed to identify members of the SAR202 clade. A total of 26 MAGs from a recent study (23) was also included in the binning process. These also were metagenomic bins from TARA metagenomes that have been assembled with megahit. The list of bins used in this study are shown in Supplemental Table 1. We also checked the bins obtained by another study using the TARA metagenomes (21) to see if there are redundant genome bins in our assemblies.

After potentially novel SAR202 bins were identified, average nucleotide identities between all TARA genome bins were determined with PyANI tool (https://github.com/widdowquinn/pyani) and a custom Python script “osu_uniquefy_TARA_bins.py” was used to identify bins that share 99% ANI. When near-identical bins were matched, more complete and less contaminated genome bin was retained. In cases where bins originated from the same TARA station, near-identical bins were combined and co-assembled with Minimus2 tool (52) to improve the genome completeness. Refinement of metagenomic bins was done using Anvi’o tool (53) to identify any potentially contaminating contigs. Some genomic bins were entirely discarded if too many multiple copies of single-copy genes are present that cannot be separated by Anvi’o. Genome completeness and redundancies were estimated using the tool CheckM (54). Genomes at various levels of completion that are less than 1.1% in redundancy of single-cope marker genes and less than 5% contamination were included for further analyses.

All the SAGs and MAGs were annotated with Prokka version 1.11 (55) to assign functions. Coding sequences predicted by Prokka were also submitted to GhostKOALA web server (56) to assign KEGG annotations to the predicted genes. In addition, Interproscan (database version 5.28-67.0) and eggNOG-Mapper (57) searches were also carried out. Metagenome-assembled genomes (MAGs) and SAGs from previous studies were also re-annotated together with the new genomes to keep the functional assignments consistent.

### Metagenome fragment recruitment analyses

Recruitment of quality-trimmed metagenomic reads from three different metagenomic databases against the SAG and MAG contigs masked to exclude ribosomal RNA-coding regions (16S, 23S, and 5S rRNA genes as predicted by barrnap) was done using FR-hit (58) with the following parameters: “-e 1e-5 -r 1 -c 80”. These parameters allowed for reads matching a given reference genome with similarity score of 80% or higher to be counted as positive matches. The metagenomic samples used for fragment recruitment were: 17 samples from BATS, 43 samples from TARA, and 22 samples from (6 from Japan, 9 from Ogasawara, and 7 from Mariana Trenches) (Supplemental Table 1). Recruitment was calculated as a percentage of quality-trimmed metagenomic reads aligned against a SAG or a MAG genome size in basepairs, normalized by total base pairs of reads in a given sample. Recruitment plot was made using “osu_plot_recruitment_heatmap.py” Python script (see https://bitbucket.org/jimmysaw/sar202_pangenomics/src/master/).

### Analysis of TARA Oceans metagenome SAR202 enzyme abundances

A custom Kraken (59) database was first built from the 122 SAR202 genomes used in this study. All coding DNA sequences in the 243 TARA Oceans metagenomic samples were then searched against the custom Kraken database containing SAR202 genomes with rRNA regions masked to identify all coding sequences belonging to SAR202 genomes.

## Data availability

All the SAGs and metagenomes are deposited to National Center for Biotechnology Information and their accession numbers are listed in the Supplemental Table 1. Prokka annotations of the genomes are available on Figshare (DOI: 10.6084/m9.figshare.8343809). All the metagenomes used for fragment recruitment analysis have been deposited to DNA Data Bank of Japan with the following submission IDs: Ogasawara Trench: DRA005790, Japan Trench: DRA005791, Mariana Trench: DRA005792. Accession numbers of each metagenomic sample are provided in the Supplemental Table 1. All code (Bash, Python, R scripts) used to analyze data and to generate figures are accessible at a Bitbucket repository (https://bitbucket.org/jimmysaw/sar202_pangenomics/src).

## Acknowledgements

We would like to thank the captain, crew, ROV and CTD operation teams, and science party of the JAMSTEC RV Kairei cruises (KR11-11, KR12-19, and KR14-01). T.N. was supported in part by a Grant-in-Aid for Scientific Research (B) (30070015) from the Japan Society for the Promotion of Science (JSPS). We thank the staff of the Bigelow Laboratory for Ocean Sciences’ Single Cell Genomics Center for the generation of single cell genomic data. We thank Mark Desenko from Center for Genome Research and Biocomputing at Oregon State University for sequencing six of the Illumina SAG libraries. The funding for mass spectrometry data collection and analysis came from the National Science Foundation (NSF Grant OCE-1154320 to EBK and KL). This work was funded by Simons Foundation International as part of BIOS-SCOPE initiative to SJG, CAC, and EBK, and by the NSF grants OCE-1335810 and DEB-1441717 to RS.

---

**Figure S1.** (**A)** Distribution of SAR202 SAGs and MAGs encoding Ring-Hydroxylating Dioxygenases (RHDs) and (**B)** SAR202-specific RHD abundances in TARA Oceans metagenomes. SAGs/MAGs with highest RHD abundances are located in coastal locations. Samples were normalized by dividing total SAR202 RHDs by total SAR202 single-copy genes found in each sample.

**Figure S2.** Maximum Likelihood phylogenetic tree of rhodopsins found in SAR202 groups based on a tree from a recent study (35). SAR202 rhodopsins are closely related to blue- and green-light absorbing proteorhodopsins (PR). Orange and white node circles indicate ultrafast bootstrap support values above and below 90, respectively.

**Figure S3.** Detailed phylogenetic tree of SAR202 rhodopsins from Figure S3, showing tips colored according to SAR202 subgroups. The phylogenetic tree was built using IQ-Tree with the following parameters: -m LG+C10+F+G -bb 1000.

**Figure S4**. Fragment recruitment of metagenomic reads from TARA Oceans metagenomic samples against all SAR202 SAGs and MAGs. Color boxes on the left of the heatmap represent different oceanic regions with the abbreviations of these oceanic regions shown in the boxes. Metagenomic samples are arranged according to depth and sample names and depth information are shown on the right of the heatmap. Branching order of the SAR202 genomes follow the order shown in the Bayesian phylogenetic tree in Figure 1.

**Figure S5**. Fragment recruitment of metagenomic reads from BATS metagenomic samples against all SAR202 SAGs and MAGs. Color boxes on the left of the heatmap represent different depths and the depth information is shown in the box. Metagenomic samples are arranged according to depth and sample names are shown on the right of the heatmap. Branching order of the SAR202 genomes follow the order shown in the Bayesian phylogenetic tree in Figure 1.

**Figure S6.** Correlation of relative enzyme abundances vs. depth of origin of most abundant paralogous families of genes in SAR202 SAGs and MAGs. The enzyme families are, **(A)** FMNOs, **(B)** enolases, **(C)** RHDs, and **(D)** dehydrogenases.

**Figure S7. (A)** Relative abundances of SAR202-specific enolases in TARA Oceans metagenome samples. Distribution of samples are plotted in order of sampling dates and depth of origin of the samples. **(B)** Correlation of normalized SAR202-specific enolase relative abundances vs. depth of origin in TARA Oceans metagenome samples. Samples were normalized by dividing total SAR202 enolases by total SAR202 single-copy genes found in each sample.

**Figure S8. (A)** World Map showing relative abundances of all FMNOs identified in all TARA Oceans metagenomes. These include SAR202-specific FMNOs and those from other organisms. Sample with highest relative abundance is highlighted in red. Different sizes of the bubbles represent the different percentages of abundance as shown in the circles below the map. **(B)** Relative abundances of FMNOs along depth profile in all TARA Oceans metagenomes. Samples are sorted in order of sampling time (from beginning to end). **(C)** Correlation between relative abundances of all FMNOs in TARA metagenomes vs. depth.

